# Acute wood smoke exposure is associated with cell-specific hippocampal transcriptomic responses in an accelerated ovarian failure mouse model

**DOI:** 10.64898/2026.01.12.699079

**Authors:** Sydnee Yazzie, Mijung Oh, Eunju Lim, Onamma Edeh, Edward Barr, Tou Yia Vue, Shuguang Leng, Eric R. Prossnitz, Robert D. Miller, Kartika Wardhani, Connor Dixson, Jennifer Gillette, Katherine E. Zychowski

## Abstract

**Background:** Wildfire events are increasing in frequency and intensity, and it is well-known that aging individuals are more susceptible to air pollution exposures, and that air pollution exposures result in neurological sequelae. Despite this, it is unclear how declining levels of ovarian hormones that naturally occur in aging females influence brain vulnerability to air pollution. Menopause and the menopausal transition represent a period of profound physiological change that affects cardiovascular, neurological, and immune health.

**Methods:** We tested whether perimenopausal–like hormonal status amplifies hippocampal responses to acute wood smoke (WS) using an ovary-intact, 4-vinylcyclohexene diepoxide (VCD) model of moderate accelerated ovarian failure (AOF) in female C57BL/6 mice. Animals were exposed to HEPA-filtered air (FA) or WS for 4 h/day over 2 consecutive days (∼0.5 mg/m³). Exposure characterization confirmed a complex mixture of combustion products with significant levels of both trace metals and gas release during WS exposure.

**Results:** Spatial transcriptomics (10x Visium; (n=4 sections/group) with automated cell-type annotation identified astrocytes, GABAergic and glutamatergic neurons, oligodendrocytes, revealed cell type-specific transcriptional alterations following WS exposure. Distinct transcriptional patterns were observed across all identified neuronal and glial cell populations.

**Conclusion:** Together, these findings define a cell type-resolved transcriptional framework linking WS exposure and ovarian hormone decline and identify potential cellular pathways relevant to hippocampal vulnerability.

## 1. Introduction

Wood smoke (WS) exposure is a growing global health concern due to an increase in both frequency and intensity of wildfires^1–4^.In recent years, hotter and drier summers, increased wind events and prolonged drought conditions have contributed to larger and more intense wildfire seasons^1–3^. With the rise of wildfire events, there is also an increase of wood smoke (WS) released in the air, exposing both rural and urban human populations to particulate matter (PM), toxic gases and unhealthy air quality^52–53,71–72^. Emissions from wildfires are naturally physically and chemically complex as it is made up of PM, gases, methane, carbon monoxide (CO), nitrous oxide (N_2_O), and trace metals^4^. These pollutants collectively pose severe risks to respiratory and cardiovascular health and are increasingly implicated in central nervous system (CNS) toxicity and neuroinflammatory responses^4–6^. Identifying biological factors that contribute to toxic susceptibility that contribute to variability in toxic susceptibility is critical for protecting populations disproportionately affected by environmental pollutants. One factor is biological sex, which has been proven to significantly influence the body’s response to air pollution in the context of immune regulation and neuroinflammation^9^. Females often exhibit heightened immune and inflammatory response in the brain which may be attributed in part to the regulatory effects of ovarian hormones on glial activation, blood-brain integrity and antioxidant systems^10–11^. Menopause is characterized by the gradual decline of ovarian hormone levels and is associated with increased systemic inflammation and could exacerbate CNS toxic susceptibility to environmental exposures^11–12^. Prior studies have demonstrated that the aging brain shows susceptibility to toxicants, including air pollution exposures^17–19,73^. Ovarian hormones naturally decline with age during the human menopausal transition, however the mechanistic contribution of this impact on neuroinflammatory sequelae following air pollution exposures is underexplored. Furthermore, the hippocampal region has demonstrated toxic susceptibility to air pollution exposures. This well-studied brain region is involved in memory, learning and neuroendocrine regulation that are particularly susceptible to neurotoxic and inflammatory effects^18,47,48^. The hippocampus is also enriched in estrogen receptors and responsive to ovarian hormone modulation, making it even more critical to understand toxicological susceptibility in an ovarian hormone deplete state^58–61^.

To address this gap in knowledge, we utilized the 4-vinylcyclohexene diepoxide (VCD), ovary-intact mouse model of accelerated ovarian failure (AOF) to explore hippocampal spatial transcriptomic alterations in an ovarian-hormone diminished mouse model of “peri-menopause.”^10,56^. This mouse AOF model has been well-established to selectively ablates primary and secondary ovarian follicles while preserving non-ovarian estrogen production and closely emulates the menopausal transition observed in women^10,12^. In comparison to surgical ovariectomy (OVX), this model offers a more physiological relevant approach to study the effects of gradual hormonal loss on neuroinflammation after acute WS exposure. This design allows for assessment of hormone-associated transcriptional responses without imposing abrupt endocrine disruption.

In this study, we utilized spatial transcriptomics to evaluate hippocampal gene expression changes following acute WS exposure in VCD-treated female C57BL/6 mice. Spatial transcriptomics allows for high-resolution gene expression profiling within intact tissue architecture^15–16^. This approach allows us to identify cell-type-specific neuroinflammatory response to acute WS exposure within a defined brain region.

We hypothesized that ovarian hormone depletion would be associated with altered hippocampal transcriptional responses to WS exposure, reflected in changes across neuronal and glial cell populations. As an exploratory analysis, these findings may offer novel insight into how ovarian hormone loss alters brain vulnerability to environmental pollutants, with implications for cell-type specific neuroinflammatory pathways and differential risk across the hippocampus.

## 2. Methods

### 2.1 Animals, and 4-vinylcyclohexene diepoxide (VCD) Injections

Six-eight week-old female C57BL/6 mice (n=8 per treatment group) were purchased from Jackson Laboratories (Ban Harbor, Maine). After a 2-week acclimation period in appropriate housing conditions, in accordance with University of New Mexico institutional animal care and use committee (IACUC) approved protocol, animals underwent an intraperitoneally (IP) at 130mg/kg for 10 days 4-vinylcyclohexene diepoxide (VCD) injection protocol^10,56^. This regime was used to induce a moderate accelerated ovarian failure state (AOF) that is physiologically relevant to human menopause state. A moderate AOF state, equivalent to peri-menopause was defined as mice that demonstrated irregular estrous cycles with approximately 6-10 days in diestrus^53–54^ (Supplemental Figure 2A).

### 2.2 Moderate-AOF Status Validation

Four-wks after injections concluded, daily vaginal cytology was conducted for 33 consecutive days prior to WS exposure to confirm moderate AOF status, characterized by an approximate 20% increase in diestrus days compared to normal cycling vehicle control females (Supplemental Figure 2A). Mice were considered to have irregular cycles and moderate-AOF after 6-10 consecutive data in diestrus^54^. Mouse plasma was collected 24-hr after exposure treatments and reduced estradiol (E2) was confirmed in the VCD-treated mice (Supplemental Figure 2B) based on a commercially available kit (ELISA Novus Biologicals, Cat # NBP3-23559).

### 2.3 Wood smoke Inhalation Protocol

Following confirmation of moderate-AOF state, analogous to human peri-menopause, mice were assigned to one of the two exposure conditions: (1) HEPA-Filtered Air (FA) or (2) acute WS exposure for 4-hr/2-d. Wood smoke-exposure was conducted by a whole-body inhalation chamber using a burned biomass wood sample (2g) sourced from local trees in the Southwestern United States. The targeted WS concentration was approximately ∼0.5mg/m^3^ +/- SEM designed to emulate an acute WS-exposure that is consistent with real-world exposure measurements (Figure 1). This system and exposure paradigm has been used in prior studies with consistent reliability and reproducibility^55–56^.

**Figure 1.**
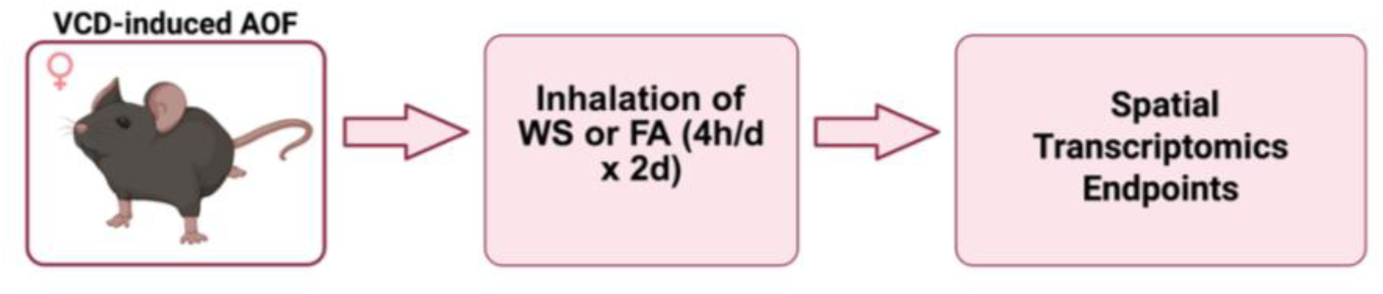
Experimental design for assessing spatial transcriptomic responses to acute wood smoke (WS) exposure. Moderate-AOF (VCD-induced ovarian failure) was assigned to one of two inhalation conditions: (a) HEPA-filtered air (FA) or (b) WS exposure (4hrs/day for 2 consecutive days). 24-hrs following last exposure, brain tissue was harvested. Spatial transcriptomic analyses was conducted to evaluate cell-type specific responses in the hippocampus.

### 2.4 Tissue Collection and Data Collection

24-hrs post-exposure, brain tissue was collected from the animals, following terminal cardiac puncture and euthanasia according to our approved IACUC protocol. The right hemisphere was embedded intact in optimal cutting temperature (OCT) compound for spatial transcriptomics using the 10X Genomics Visium platform, focusing on the hippocampus to capture cell-type-specific gene expression data for astrocytes, microglia and other cell types^74^.

### 2.5 Tissue Slicing and RNA Extraction Preparation

Slicing and hematoxylin and eosin (H&E) staining was performed by the University of New Mexico Human Tissue Repository (HTR). Tissue sections were prepared at a thickness of 5μm for H&E staining. The hippocampal region was identified and selected as the area of interest for spatial transcriptomic analysis and 4 selected areas ∼2mm in diameter were from the hippocampal region and included the CA3, CA1-CA3 and CA3-dentate gyrus junctions for downstream spatial transcriptomics sequencing. Given the 2 mm diameter of Visium sequencing spots, this represents the maximal spatial specificity achievable for region-of-interest (ROI) selection^82–84^.

Additional tissue sections were prepared at a thickness of 5μm and mounted onto slides designated for RNA extraction and quality control (QC), 2 slides per sample per exposure groups. Following sectioning, slides were immediately stored at -80 °C to preserve RNA integrity. Slides were then shipped on dry ice to Northwestern University Center for Genetic Medicine in Chicago, IL. After arrival, tissue sections were manually scraped from the slides, and total RNA was extracted. As section thickness does not impact RNA extraction efficiency in this procedure, minor variations in thickness were negligible. RNA integrity was assessed post-extracted, and results were communicated with the Zychowski lab. After RNA QC was determined acceptable, additional tissue sections were prepared for spatial transcriptomics.

In the second set of sample preparation, two tissue sections per sample were mounted onto slides, with a required thickness of 10μm. Unlike formalin-fixed and paraffin-embedded (FFPE) tissues, no incubation or drying steps were performed. Immediately after sectioning, slides were stored at -80 °C to preserve tissue integrity, Slides were shipped on dry ice to Northwestern University Center for Genetic Medicine in Chicago, IL for further processing. Upon arrival, one slide per sample, was selected for the Visium CytAssist experiment, which was conducted within two weeks of sample arrival.

### 2.6 Bioinformatics Analysis

Spatial transcriptomic libraries were prepared using the Visium Spatial Gene Expression platform (10x Genomics, Pleasanton, CA, USA). Tissue sections were imaged using immunofluorescence and the spatially barcoded transcriptomics were captured. Image processing, spot detection and alignment were performed done using Space Ranger (10X Genomics, Pleasanton, CA, USA), which mapped spatial barcodes to their respective locations on the tissue sections. Quality control metrics, including the number of unique molecular identifiers (UMIs), mitochondrial read content and spatial barcode detection efficiency were assessed. Spots with low RNA content or high mitochondrial gene expression were removed from downstream analysis. Two mice per group, each with two brain sections were analyzed (n = 4 sections per treatment group). Four spots from each hippocampal slice (n = 4 x 4 = 16 spots total) were selected within the hippocampus from both FA and WS-exposed animals for comparative analysis (Supplementary Figure 1). The study design follows precedents in neurological and toxicological mouse studies with similar replication schemes, demonstrating the feasibility and reproducibility of such designs for high-content spatial transcriptomic analyses^78–82^.

FASTQ files were processed using Space Ranger, which included adaptor trimming, alignment trimming, alignment to the reference genome using the STAR aligner, and assignment of spatial barcodes to their corresponding transcriptomic reads. A gene expression matrix was generation, associating detected transcripts with individual spatial spots.

Gene expression values were normalized to correct for sequencing depth and technical variability. Dimensionality reduction was done, followed by unsupervised clustering and spatially variable genes were identified by spatial autocorrelation methods.

To identify cell types within the spatially resolved transcriptome, scType (Ianevski et al., https://github.com/IanevskiAleksandr/sc-type), an automated cell type predicated algorithm was implemented. scType uses a database of known marker genes for various cell types for different tissues. The software computes a likelihood score for each cell and determines the probability that it belongs to that particular cell types based on its gene expression profile. Cells with high confidence scores were assigned a specific cell type, while those with insufficient confidences were categorized as “Unknown.” Markers from scType for brain cell types including, astrocytes, cholinergic neurons, endothelial cells, GABAergic neurons, glutamatergic neurons, microglia, oligodendrocyte progenitor cells (OPCs) and oligodendrocytes, were crossed referenced in Brain RNA-seq (https://brainrnaseq.org) to confirm their specificity to its respective cell type. This approach allows for accurate and valid cell type annotation for spatial transcriptomic identification of hippocampal-specific gene expression patterns^75,76^.

### 2.7 Data Visualization

Downstream analysis in R Studios (R Core Team, 2025) included violin plot generation to visualize gene expression distribution in our identified cell types in the hippocampus. UMAPS were created to visualize transcriptional clustering of cell types and highlight cell-type specific expression patterns in our region of interest. To further interpret the biological significance, differentially expressed genes were identified between spatially distinct regions by DESeq2, followed by pathway enrichment analysis using Gene Ontology (GO), Kyoto Encyclopedia of Genes and Genomes (KEGG). Spatial transcriptomic data were visualized using Seurat’s SpatialFeaturePLot and Scanpy’s spatial plotting functions, overlaying gene expression patterns on the tissue section image. Clustered regions and spatially enriched genes expressions profiles were mapped using ggplot2 for high-resolution spatial feature visualization. This workflow provides a comprehensive approach to analyzing 10X Genomics Visium spatial transcriptomics data, from raw sequencing reads to biologically relevant spatial information^77^.

## 3. Results

### 3.1 Metal and Gas Composition During WS Exposure

Analysis of particulate matter (PM) metals collected from FA and WS filters (n=6 per group) demonstrated a significant increase of metals in WS-exposed samples during whole-body inhalation exposure. Specifically, ICP-MS analysis on isolated filter samples identified a statistically significant increase in copper (^63^Cu) (p=0.0320) and nickel (^60^Ni) (p=0.0444) in WS filters compared to FA controls (Figure 2 A-B), consistent with metal emissions from biomass combustion during real-world wildfire events. While other trace metals including tungsten, lead, uranium, sodium, magnesium, aluminum, potassium, vanadium, chromium, manganese, iron, cobalt, zinc, arsenic, selenium, cadmium, calcium, antimony, barium, thallium, and thorium did not show statistically significant changes in our samples.

**Figure 2.**
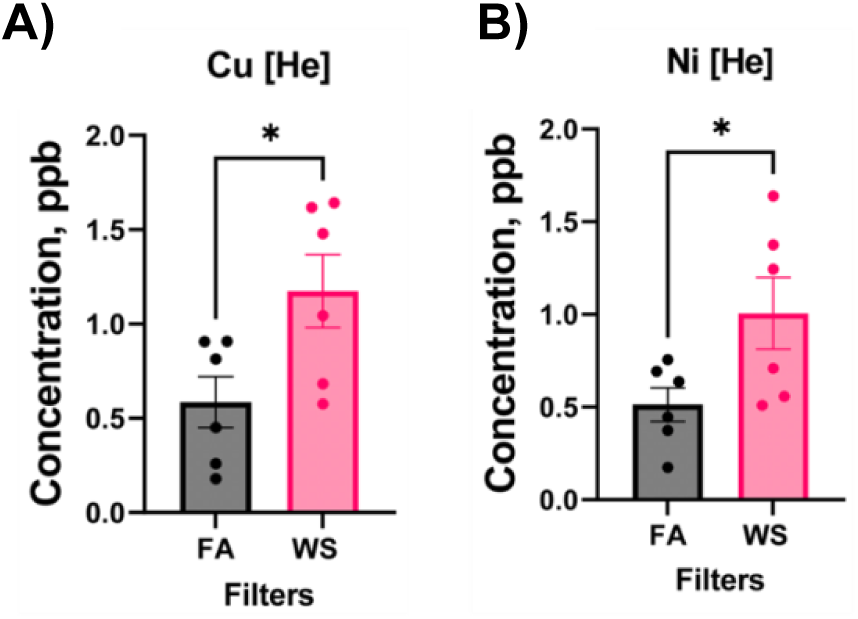
ICP-MS Metals Analysis in FA and WS filter samples. Metals analysis revealed a significant increase (p<0.05) of Cu (A) and Ni (B) in WS filters collected after exposure period, compared to FA (n=6/ treatment group).

In parallel, real-time monitoring of gas release during WS exposure demonstrated variable patterns in combustion-related pollutants throughout the exposure period (Figure 3 A-E). Sulfur dioxide (SO_2_) and carbon dioxide (CO_2_) levels gradually increased while nitrogen dioxide (NO_2_) demonstrated episodic spikes. Carbon monoxide (CO) showed transient peaks while Ozone (O_3_) release remained relatively low but had brief surges. These findings confirm that the WS exposure chamber environment contained complex mixtures of inhaled metals and reactive gases consistent with real world biomass combustion emissions.

**Figure 3.**
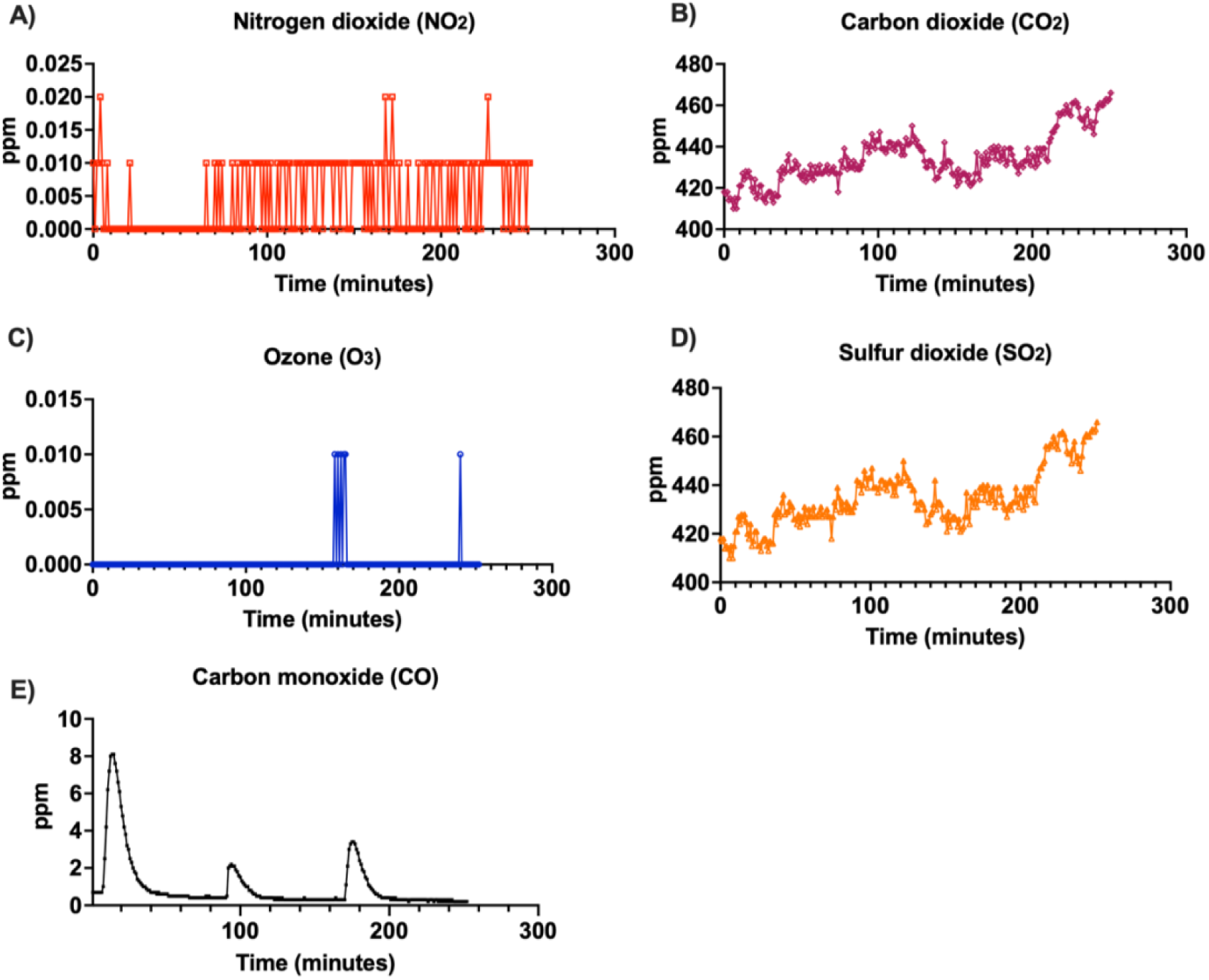
Wood smoke gases analysis. Gases sampled during WS exposure including real-time measurements of A) nitrogen dioxide (NO_2_), B) carbon dioxide (CO_2_), C) ozone (O_3_), D) sulfur dioxide (SO_2_), and E) carbon monoxide (CO)

### 3.2 Hippocampal Cell Type Clustering and Distribution Following Acute WS Exposure in a Moderate-AOF Mouse Model

UMAP projections identified distinct clustering of key hippocampal cell populations including astrocytes, GABAergic neurons, glutamatergic neurons, oligodendrocytes and oligodendrocyte precursor cells (OPCs) (Figure 4A).

**Figure 4.**
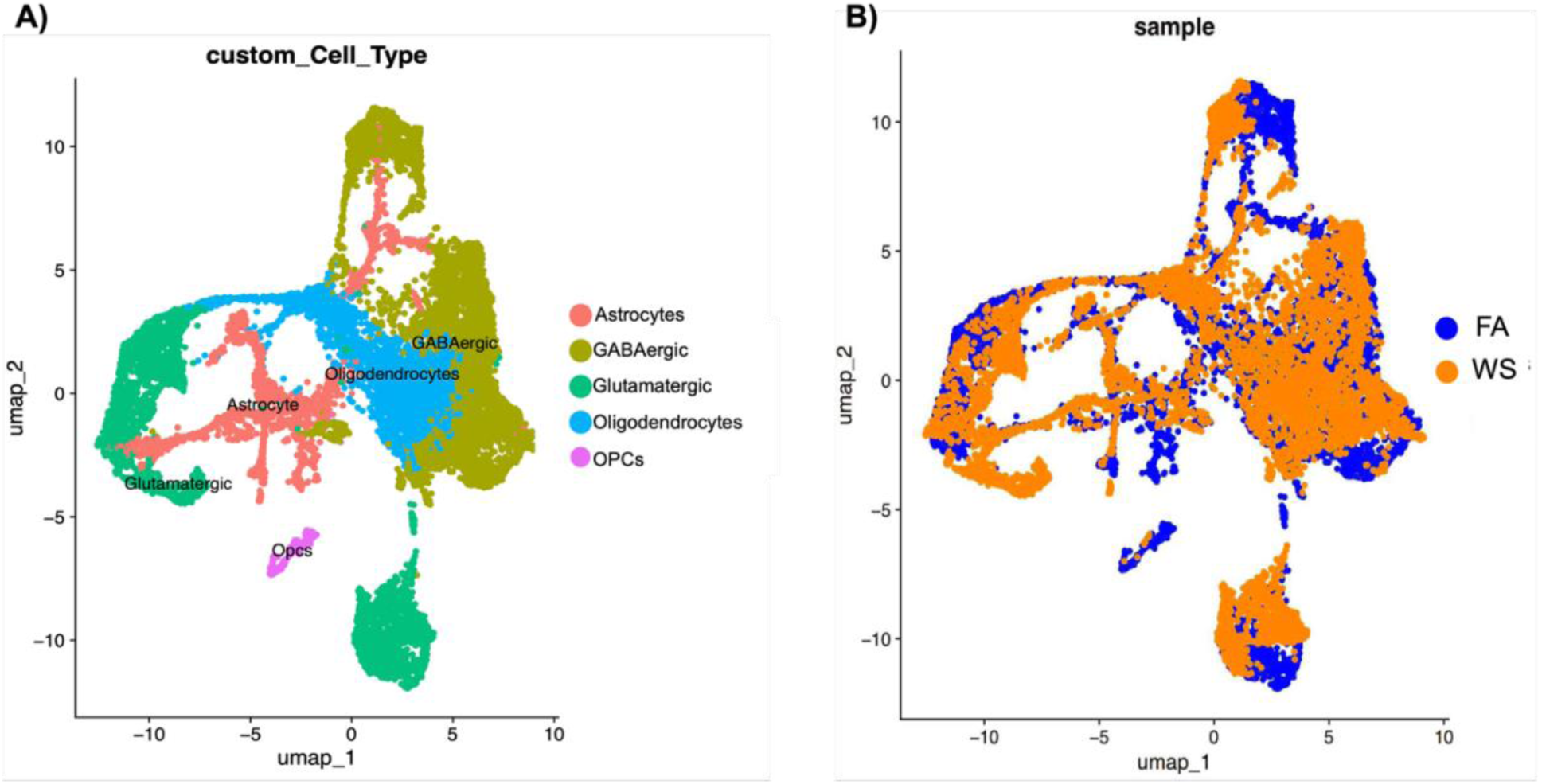
UMAP visualization of hippocampal cell type distribution according to each treatment. Cell type UMAP (A) shows distinct cell type clustering of astrocytes, GABAergic neurons, glutamatergic neurons, oligodendrocytes and oligodendrocyte precursor cells (OPCs). Exposure conditions UMAP (B) demonstrates distribution of cell types under experimental conditions FA (blue) and WS (orange).

When separated based on exposure, apparent shifts in the density and representation of select cell types between FA and WS suggest that wood smoke exposure influences transcriptional states within identified cell populations (Figure 4B). This observation is consistent with prior work demonstrating that hippocampal cellular composition and gene expression can be dynamically modulated by age, injury, or genetic factors^18–19,49^.

### 3.3 Distinct Transcriptional Alterations in GABAergic and Glutamatergic Neurons Following WS Exposure

To determine whether WS exposure influences transcriptional responses between excitatory and inhibitory neurons within the hippocampus, we compared differentially expressed gene profiles of GABAergic and glutamatergic neuronal populations. GABAergic neurons demonstrated transcriptional activation of genes associated with transcriptional/developmental regulation (*Sp8*, *Prdm12*, *Zic4*), immune or chemokine-related signaling (*Prkcd*), and neuroendocrine modulation (*Pnoc*). (Figure 5A).

**Figure 5.**
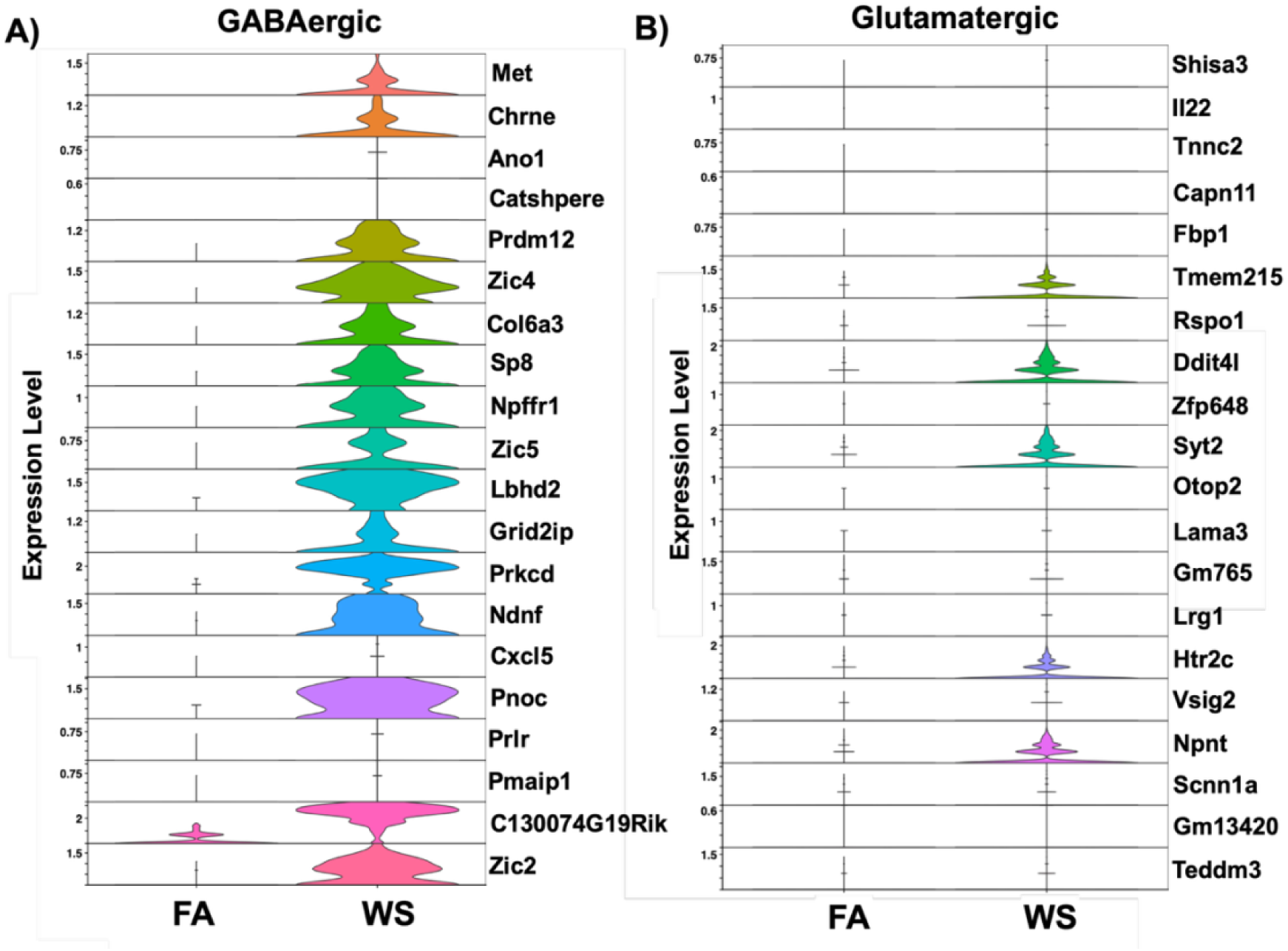
Violin plots demonstrating key gene expression levels associated with hippocampal neurons. A) GABAergic neurons, and B) glutamatergic neurons. Visualized differential gene expression levels were based on a significant p-value of ≤0.05 and a log-fold change of 1. Each plot illustrates variability and distribution of gene expression in different hippocampal cell types under FA and WS exposure conditions (n=4 sections/group).

Glutamatergic circuits displayed stress-adaptive transcriptional shifts highlighted by *Ddit4l*, a stress-inducible mTOR inhibitor that promotes autophagy and metabolic restraint under hypoxic/oxidative challenge, consistent with cellular stress-adaptation programs observed *in vivo* and in primary cells^20,23^. Postsynaptically, *Htr2c* signaling enhances hippocampal plasticity and memory, providing a serotonergic route by which stress and metabolic states can tune excitatory circuits^24^ (Figure 5B).

### 3.4 Oligodendrocyte and Astrocyte-Specific Transcriptional Responses Following WS Exposure

In addition to neuronal populations, we examined transcriptional alterations in oligodendrocytes and astrocytes, two glial populations critically involved in maintaining hippocampal circuit stability.

Astrocytes exhibited a more complex transcriptional response. WS exposure led to reduced expression of stress-response and chromatin remodeling transcripts, such as *Hspa1b* and *Junb*, while upregulating synaptic regulation and glutamate transporter-associated genes including *Syt2*, *Slc17a6*, and *Amigo2* (Figure 6A). Oligodendrocytes displayed increased expression of genes associated with developmental regulation and synaptic adaptation, including *Zic1*, *Tcf7l2*, and *C1ql2* (Figure 6B).

**Figure 6.**
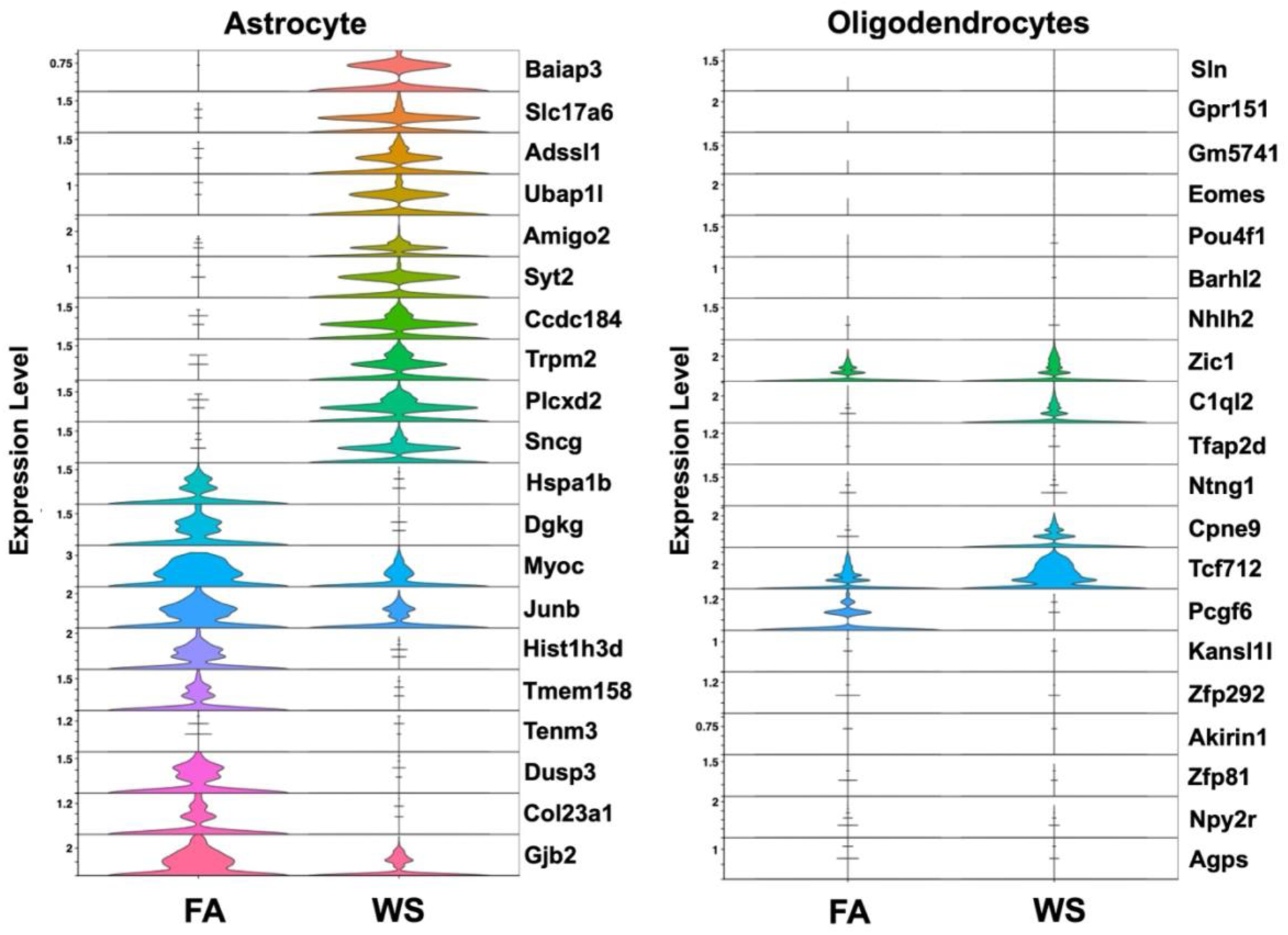
Violin plots demonstrating key gene associated with hippocampal non-neuronal cells. A) astrocytes, B) oligodendrocytes. Visualized differential gene expression levels were based on a significant p-value of ≤0.05 and a log-fold change of 1. Each plot illustrates variability and distribution of gene expression in different hippocampal cell types under FA and WS exposure conditions (n=4 sections/group).

### 3.5 Enriched Gene Ontology Biological Processes in GABAergic and Glutamatergic Neurons Following WS Exposure

In GABAergic neurons, upregulated enrichment was observed in pathways related to synaptic signaling and neurotransmission, including modulation of chemical synaptic transmission (GO:0050804) and synapse organization (GO:0050808). Additional categories reflected broad neurodevelopmental and plasticity-associated processes, such as neuron projection development (GO:0031175), glial cell differentiation (GO:0010001), and regulation of neuronal synaptic plasticity (GO:0048168) (Figure 7A).

**Figure 7.**
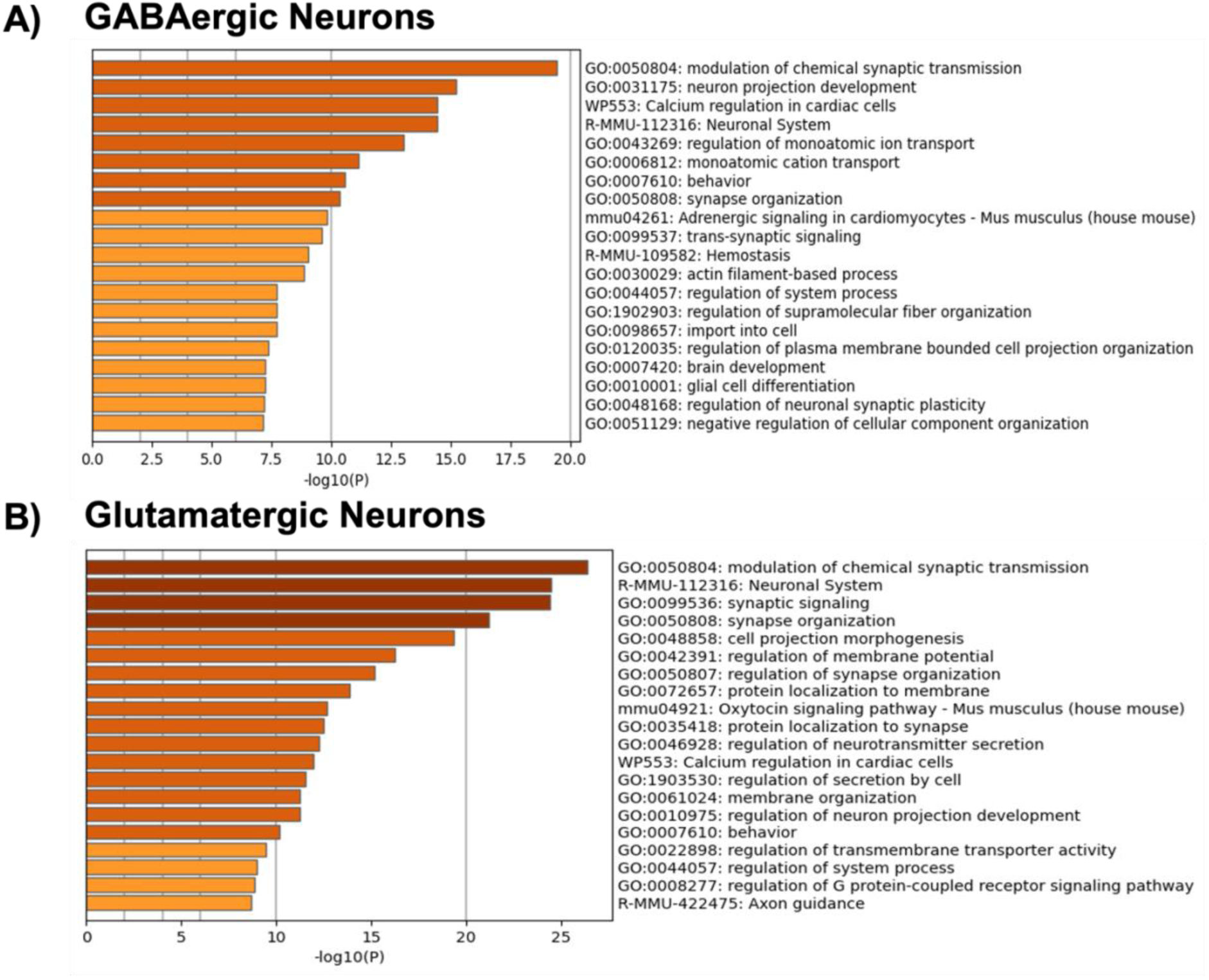
Upregulated enriched ontology clusters in hippocampal neurons in WS compared to FA. GO/KEGG pathway analysis of A) GABAergic and B) glutamatergic neurons in WS compared to FA exposure. Enriched terms, identified through statistical enrichment analysis, were grouped into clusters based on gene membership similarities

In glutamatergic neurons, enrichment was more pronounced in pathways regulating excitatory transmission and structural remodeling, including modulation of chemical synaptic transmission (GO:0050804), axon guidance (R-MMU-422475), and cell projection morphogenesis (GO:0048858). Upregulation of protein localization to synapse (GO:0035418) and regulation of synapse organization (GO:0050807) categories highlight the vulnerability of excitatory circuits to metabolic and oxidative stress (Figure 7B), which may underlie hyperexcitability phenotypes observed following particulate exposure^17^.

### 3.6 Enriched Gene Ontology Pathways in Oligodendrocytes and Astrocytes Following WS Exposure

In astrocytes, upregulated enriched pathways included neurotransmitter transport (GO:0051588), cell morphogenesis involved in neuron differentiation (GO:0048667), and central nervous system myelination (GO:0022010) (Figure 8A).

**Figure 8.**
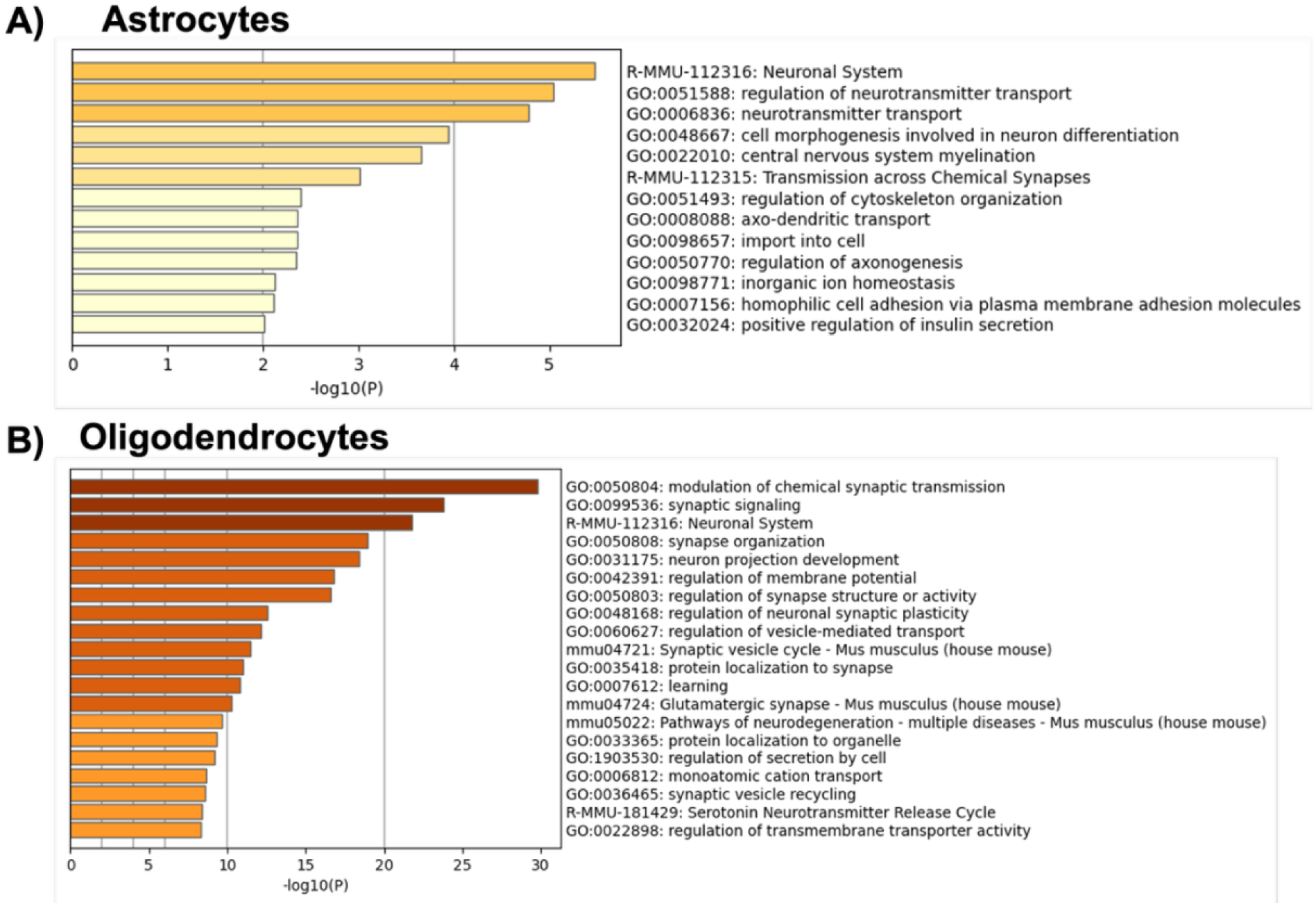
Upregulated enriched ontology clusters in hippocampal non-neuronal cells in WS compared to FA. GO/KEGG pathway analysis of A) astrocytes, and B) oligodendrocytes. Enriched terms, identified through statistical enrichment analysis, were grouped into clusters based on gene membership similarities

In oligodendrocytes, enrichment was broader and strongly linked to synaptic and metabolic regulation, including modulation of chemical synaptic transmission (GO:0050804), synaptic signaling (GO:0099536), synapse organization (GO:0050808), and regulation of neuronal synaptic plasticity (GO:0048168). Additional categories included serotonin neurotransmitter release cycle (R-MMU-181429) and pathways of neurodegeneration (mmu05022) (Figure 8B).

### 3.7 Downregulated Gene Ontology Pathways in Neuronal Cell Populations following WS Exposure

In GABAergic neurons, significant downregulation was observed in pathways central to neuronal communication and learning, including behavior (GO:0007610), synaptic signaling (GO:0099536), regulation of membrane potential (GO:0042391), and learning (GO:0007612). Several categories related to ion transport and signaling cascades were also suppressed, such as regulation of monoatomic ion transport (GO:0043269), G protein–coupled receptor signaling (R-MMU-372790), and dopaminergic synapse (mmu04728) (Figure 9A).

**Figure 9.**
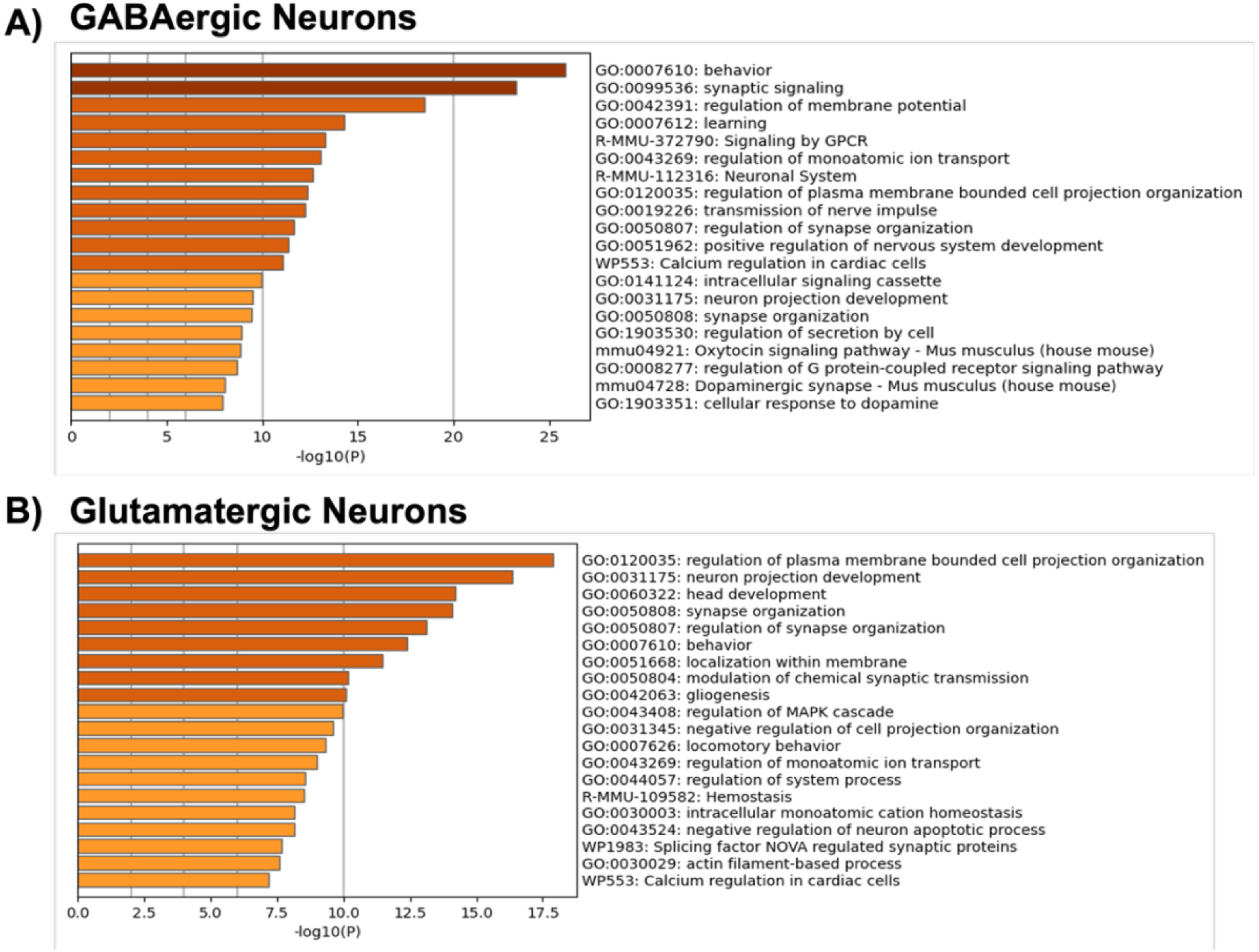
Downregulated enriched ontology clusters in hippocampal neuronal cells in WS compared to FA. GO/KEGG pathway analysis of A) GABAergic, and B) glutamatergic neurons. Enriched terms, identified through statistical enrichment analysis, were grouped into clusters based on gene membership similarities

In glutamatergic neurons downregulation extended to developmental and structural pathways, including neuron projection development (GO:0031175), head development (GO:0060322), and gliogenesis (GO:0042063). Suppression of MAPK cascade regulation (GO:0043408) and negative regulation of apoptotic processes (GO:0043524) suggests impaired pro-survival signaling and stress response regulation (Figure 9B).

### 3.8 Downregulated Gene Ontology Biological Processes in Non-neuronal Cell Populations following WS Exposure

In astrocytes, we identified downregulation of pathways related to tissue structure and repair, including collagen biosynthesis and modifying enzymes, ossification (R-MMU-1650814), connective tissue development (GO:0061448), and glial cell proliferation (GO:0014009). Additional suppression of MAPK signaling (mmu04010), response to peptide hormone (GO:0043434), and prostaglandin biosynthetic processes (GO:0001516) (Figure 10A)

**Figure 10.**
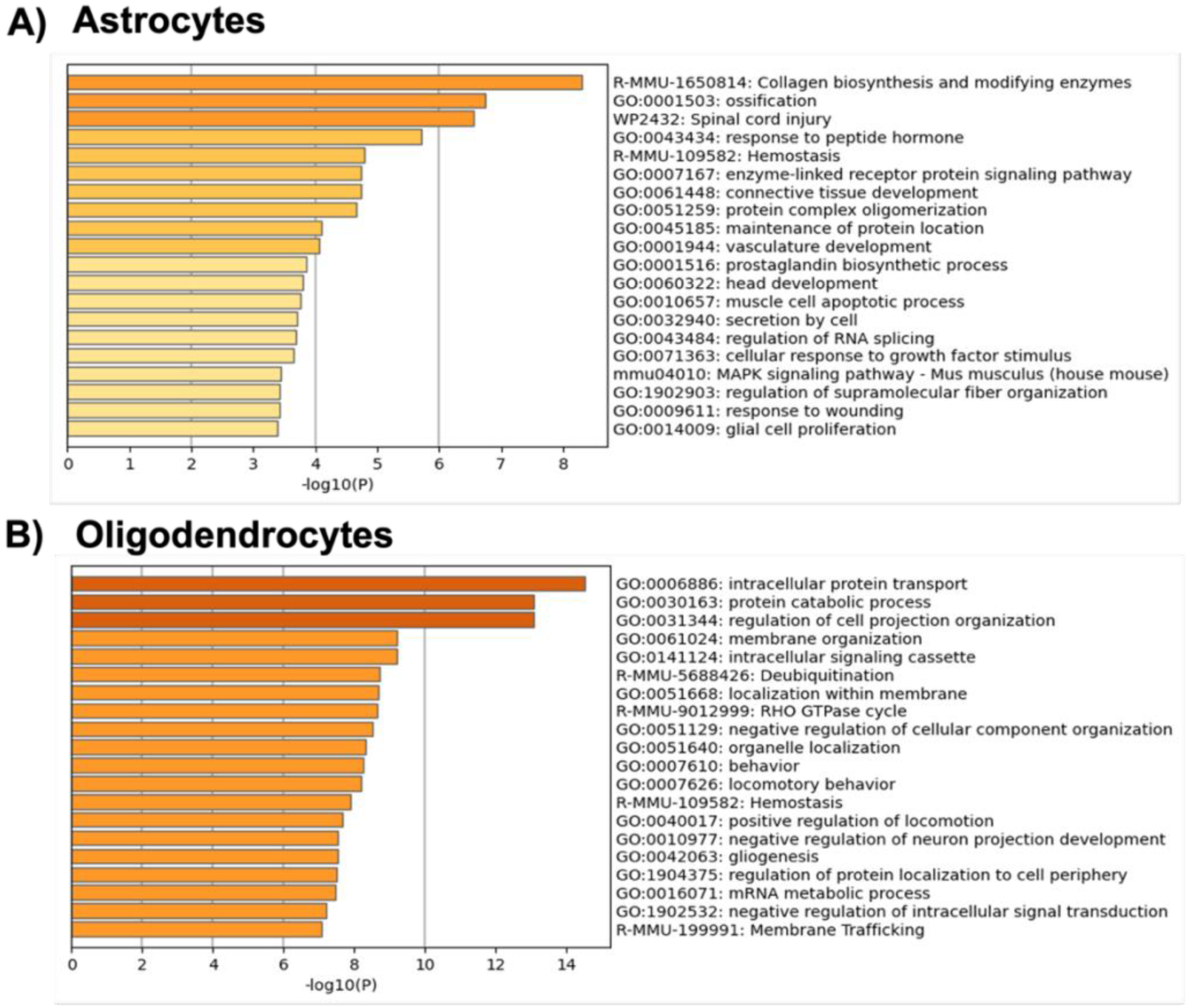
Downregulated enriched ontology clusters in hippocampal non-neuronal cells in WS compared to FA. GO/KEGG pathway analysis of A) astrocytes, and B) oligodendrocytes. Enriched terms, identified through statistical enrichment analysis, were grouped into clusters based on gene membership similarities

In oligodendrocytes, we observed that downregulated categories were linked to intracellular transport and signaling, such as protein catabolic process (GO:0030163), intracellular signaling cassette (GO:0141124), RHO GTPase cycle (R-MMU-9012999), and membrane trafficking (R-MMU-199991). Suppression of gliogenesis (GO:0042063), regulation of cell projection organization (GO:0031344), and mRNA metabolic process (GO:0016071) (Figure 10B).

## 4. Discussion

This exploratory study provides novel insights into hippocampal transcriptional responses following acute WS exposure in a physiologically relevant model of menopause-like ovarian hormone decline. Using the 4-vinylcyclohexene diepoxide (VCD)-induced model of accelerated ovarian failure (AOF), we characterized hippocampal transcriptional alterations following acute WS exposure containing both gaseous and metals trace elements. Given the well-established sensitivity of the hippocampus to hormonal fluctuations and inflammatory perturbations^5,9,58,64^, it is an ideal region of interest to evaluate cell-type specific responses to inhaled environmental toxicants in an ovarian hormone depleted mouse model, especially based on our prior findings^58,59,85^.

We confirmed that the WS in our whole-body exposure chamber contained complex mixtures of both traces metals such as ^63^Cu and ^60^Ni and release of combustion-related gases such as sulfur dioxide (SO_2_), nitrogen dioxide (NO_2_), carbon dioxide (CO_2_), carbon monoxide (CO), and ozone (O_3_) throughout the 4-hr/2d exposure period (Figure 2-3). This exposure profile reflects the chemical complexity of biomass combustion and is consistent with reported characteristics of acute wildfire smoke events^4,65,70^. Metals analysis revealed significant enrichment of ^63^Cu and ^60^Ni in WS filters compared to filtered air (FA) filters, both of which are well-documented components of wildfire particulate matter (PM) and have been associated with pro-inflammatory and neurotoxic signaling in other studies^3,4,65,70^.Concurrently, we confirmed that WS in our whole-body exposure chamber contained trace combustion gases including SO₂, NO₂, CO₂, CO, and O₃. Concentrations were elevated relative to FA baselines but generally fell within or slightly above commonly cited urban background levels. Importantly, our measured gas levels were substantially lower than many reported wildfire plume peaks^65,67,70^. For example, airborne campaigns in the western U.S. have documented CO enhancements reaching tens to hundreds of ppm and NOₓ increases of tens to hundreds of ppb in fresh plumes^65,63^. Reviews of wildfire-associated ozone chemistry similarly report O₃ enhancements of tens of ppb above background, with some events producing >100 ppb in impacted regions^67,68^. More recent field datasets from FIREX-AQ and other campaigns confirm enhancement ratios for trace gases that substantially exceed ambient levels, consistent with plume-scale emissions^70^. Regionally, O_3_ exceptional event analyses also indicate wildfire contributions of ∼19 ppb to daily maximum 8-hr ozone in western U.S. cities^69,72^. Therefore, while gases were not the predominant component of our exposure environment compared to PM and inorganic metals, they remain plausible contributors to oxidative and inflammatory stress under a mixed-exposure framework. This exposure profile underscores the relevance and significance of our exposure protocol for investigating acute WS induced neurotoxicity and its intersection with female gonadal hormonal status.

Spatial transcriptomics allows for evaluation of transcriptional alterations at single-cell resolution within the hippocampus^15–16^. Using scType annotation and high-confidence marker gene sets, we identified distinct neuronal and glial cell populations exhibiting WS-associated transcriptional changes, including astrocytes, oligodendrocytes, and neuronal subtypes. Early transcriptomic patterns suggest that under an ovarian hormone diminished state WS-induced neuroinflammatory signaling is upregulated, aligning with prior studies implication estradiol in mediating glial reactivity, and oxidative stress responses^27,40,58–59^. This study provides novel insights into the molecular and cell type–specific responses of the hippocampus, a key region for learning and memory processing, to acute WS exposure in the context of ovarian hormone decline, using spatial transcriptomics to interrogate neuronal and glial populations. Our findings demonstrate that both excitatory and inhibitory neurons, as well as astrocytes and oligodendrocytes, exhibit distinct transcriptional alterations following WS exposure, within an ovarian hormone deplete state. This dual influence of environmental toxicants and ovarian hormone decline highlights a critical intersection of environmental and biological risk factors for hippocampal vulnerability.

Analysis of neuronal populations revealed divergent transcriptional changes between inhibitory and excitatory neurons. GABAergic neurons exhibited enrichment of transcriptional regulators (*Sp8, Prdm12, Zic4*), immune signaling genes (*Prkcd,*), consistent with potential compensatory inhibitory plasticity aimed at maintaining network stability under stress. In contrast, glutamatergic neurons exhibited upregulation of excitatory signaling and remodeling transcripts (*Tmem215, Ddit4l*), aligning with prior studies showing heightened glutamatergic sensitivity to oxidative and metabolic stress^42,44^. While these transcriptional profiles do not establish functional outcomes, they do suggest differential engagement of inhibitory and excitatory neuronal programs following WS exposure, consistent with other studies of particulate matter induced neurotoxicity^26–28,39^.

In glial populations, oligodendrocytes showed enrichment of adaptive myelination and morphogenetic programs (*Zic1, C1ql2*), consistent with adaptive remodeling to maintain myelin integrity induced by toxicant stess^32,33^. Estradiol and ERβ signaling normally promotes both lipid and cholesterol biosynthesis needed for myelin maintenance and ovarian hormone decline may alter the balance of reparative mechanisms increasing reliance on compensatory pathways^35^. Astrocytes demonstrated a dual pattern of suppressed stress-response transcripts (*Hspa1b, Junb*) and enriched glutamate transporter/synaptic support genes (*Slc17a6, Amigo2*). These findings are consistent with evidence that glial cells dynamically remodel in response to environmental stressors, with oligodendrocytes showing activity-dependent plasticity^32^ and astrocytes exhibiting both protective and pro-inflammatory phenotypes^31^. Importantly, enrichment of neurodegeneration-related pathways in astrocytes suggests that acute exposures may initiate early molecular changes that warrant further investigation for potential long-term relevance. Reduced estrogenic support has been associated with impaired glutamate uptake, oxidative stress and increased neuroinflammatory vulnerability in cortical-hippocampal circuits^36,38,45^, consistent with the synaptic remodeling and neurodegeneration-related pathways enrichment observed here^17^.

Estrogen signaling through ERα/ERβ/GPER supports synaptic plasticity, glutamate clearance, and myelin maintence^47–48^. In this study, transcriptional patterns associated with reduced MAPK/PI3K-Akt signaling, gliogenesis and stress-response pathways were observed in the context of ovarian hormone decline. These results parallel clinical imaging findings that menopause accelerates hippocampal vulnerability to metabolic and inflammatory stress^46,48^ and experimental studies demonstrating that estrogenic support preserves dendritic spines and cognitive resilience^42^. Collectively, our transcriptional findings support a working hypothesis that ovarian hormone decline and WS exposure interact to shape hippocampal molecular response, identifying potential pathways and cellular processes for further mechanistic and functionally validated investigations.

A few modest but noted limitations should be mentioned, regarding this study. First, our analysis was restricted to acute WS exposure in a moderate VCD-induced accelerated ovarian failure model, which emulate key features of the human peri-menopause transition, therefore, relationships between chronic exposures, long-term hormone decline, and neuroinflammatory outcomes remain to be fully elucidated. While chronic exposures and later stages of hormone decline were not included in the current study, the selected model and exposure paradigm were intentionally designed to preliminary isolate early hormone-dependent neuroimmune and transcriptomic responses to WS. This design supports robust identification of exposure and hormone sensitive pathways and provides a strong foundation for hypothesis generation and targeted mechanistic follow-up in future studies. Furthermore, a key next step will be protein-level validation of oxidative stress and inflammatory signaling markers such as 8-oxo-dG and NF-kB, alongside glial and synaptic makers including GFAP, S100B and synaptophysin using targeted ELISAs and/or proteomic approaches. This analysis will further clarify how the transcriptional alterations observed here translate into pathological outcomes. Lastly, although our findings align with clinical studies linking menopause to brain vulnerability^53–54^, translational work directly examining peri- and postmenopausal women in wildfire smoke exposed populations is needed. Future studies should incorporate chronic exposure paradigms, longitudinal hormone replacement strategies, and multimodal approaches (transcriptomics, proteomics, imaging) to more comprehensively define mechanisms of interaction. This pilot spatial-transcriptomics study analyzed multiple brain sections per treatment group, enabling within-animal replication and revealing consistent spatial patterns across sections. However, due to the exploratory and cross-sectional nature of our study, further studies will be needed to extend these findings to firmly establish causality.

Taken together, this study highlights significant cell type-specific and hormone-sensitive transcriptional responses in the hippocampus following WS exposure. By identifying transcriptional programs linked to excitatory-inhibitory balance, glial reparative capacity, and hormone regulation, provides a focused framework for future mechanistic and translational investigation. These findings establish a molecular reference for prioritizing cellular targets and signaling pathways relevant to wildfire smoke exposure, with particular relevance to peri- and postmenopausal populations.

## 5. Conclusions

Together, these data define broad, cell type–specific transcriptional adaptations across the hippocampus in response to acute WS exposure. Neurons exhibited divergent responses, with GABAergic populations showing enrichment of stress-responsive and plasticity-associated programs, while glutamatergic neurons displayed increased expression of genes related to excitatory signaling and synaptic remodeling pathways. Among glial cells, oligodendrocytes were characterized by transcriptional shifts in developmental and myelination-related pathways, whereas astrocytes displayed a dual profile of reduced stress-response signaling coupled with enhanced synaptic regulation and glutamate transport. Across all identified cell types, functional enrichment analyses highlighted convergence on pathways involved in synaptic reorganization, neurotransmission, and structural remodeling, alongside relative suppression of developmental and reparative programs. When considered in the context of ovarian hormone decline, these transcriptional patterns suggest increased vulnerability of hippocampal circuits, potentially reflecting altered MAPK/P13K-Akt signaling, reduced glial support capacity and changes in synaptic stability. Importantly, these findings represent an exploratory, transcriptomic framework that identifies candidate cellular processes and signaling pathways sensitive to the combined effects of wood smoke exposure and hormonal status. Future studies incorporating broader scale and functional outcomes may be conducted in the future to determine how these transcriptional alterations translate into neuroinflammatory outcomes and cognitive or behavioral relevance.

## CRediT authorship contribution statement

Conceptualization, S.Y., K.E.Z.; methodology, S.Y, T.Y.V. investigation S.Y., M.O., E.L., O.E., E.B., K.W., C.D.; data curation, S.Y., M.O., E.L., S.L.; writing-original draft, S.L., R.M., J.G., K.E.Z., ; writing-review & editing, S.Y., J.G., K.E.Z, ; funding acquisition K.E.Z.;, resources, E.P., K.E.Z.; supervision, K.E.Z

## Funding

This work was supported by the National Institutes of Health (NIH) grants: R01 ES0339881, NM-INSPIRES P30ES032755 and IMSD T32 GM144834. Additional resource support was provided from the Autophagy Inflammation and Metabolism (AIM) Center (P20FM121176) and the UNM Comprehensive Cancer Center (UNM-CCC) Shared Resources (P30CA118100).

## Data Availability

The sequencing data reported in this manuscript has been deposited into the Dryad digital repository.

## Conflicts of Interest

The authors declare no competing interests.

## Acknowledgements

The authors would like to thank Dr. Matthew J. Campen and the NM-INSPIRES Translational Research Support Core (TRSC) for usage of the inhalation system for the WS exposures. This work was supported by the Northwestern University NUSeq Core Facility.

---

**Supplemental Figure 1.**
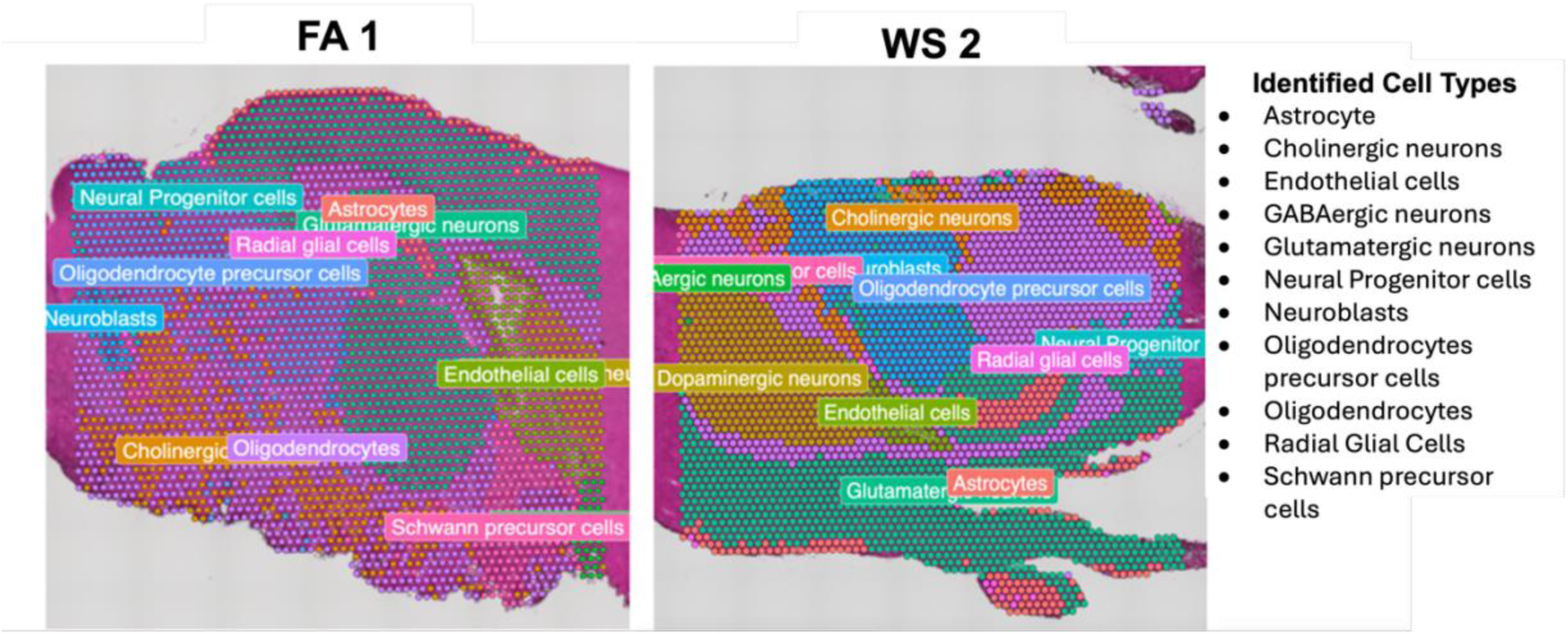
Spatial transcriptomic analysis highlights distinct cell population distribution within female mouse hippocampi samples under FA and WS exposure conditions (n=4 sections/group).

**Supplemental Figure 2.**
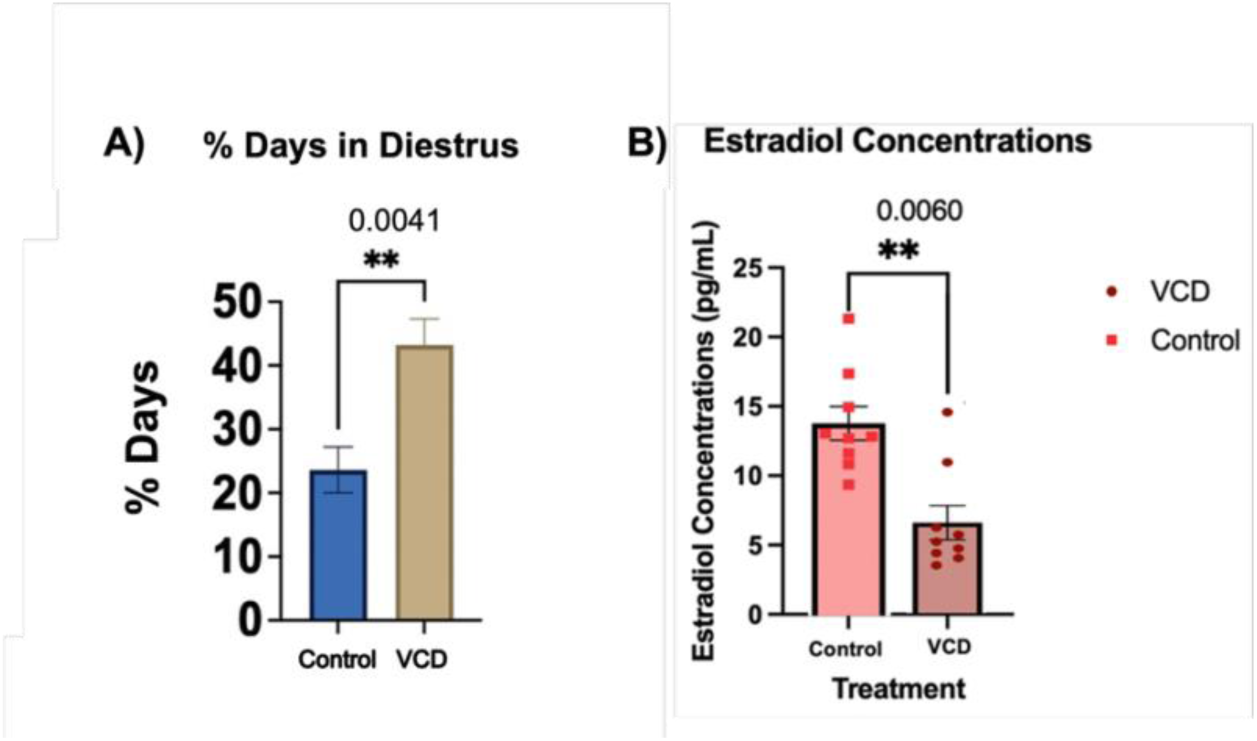
Validation of moderate-AOF state in female C57BL/6 mice. (A) Percentage of days spent in diestrus for control and moderate-AOF VCD treated animals. Daily Vaginal cytology was conducted to track estrous cycles. Statistical significance is defined as p≤ 0.05. **p≤0.01. (B) ELISA analysis of plasma estradiol concentrations (pg/mL) in control (sesame oil-injected) and moderate-AOF VCD animals (n=9/per group).

## Notes

### Competing Interest Statement

The authors have declared no competing interest.

